# Label-free high-resolution 3-D imaging of gold nanoparticles inside live cells using optical diffraction tomography

**DOI:** 10.1101/097113

**Authors:** Doyeon Kim, Nuri Oh, Kyoohyun Kim, SangYun Lee, Chan-Gi Pack, Ji-Ho Park, YongKeun Park

**Affiliations:** Department of Physics, Korea Advanced Institute of Science and Technology (KAIST), 291, Daehak-ro, Yuseong-Gu, Daejeon 34141, Republic of Korea; Tomocube, Inc., 48, Yuseong-daero 1184beon-gil, Yuseong-Gu, Daejeon 34051, Republic of Korea; Department of Bio and Brain Engineering, KAIST, 291, Daehak-ro, Yuseong-Gu, Daejeon 34141, Republic of Korea; KAIST Institute for Health Science and Technology (KIHST), KAIST, 291, Daehak-ro, Yuseong-Gu, Daejeon 34141, Republic of Korea; Asan Institute for Life Sciences, University of Ulsan College of Medicine, 88, Olympic-ro 43-gil, Songpa-Gu, Seoul 05505, Republic of Korea

**Keywords:** label-free imaging, optical diffraction tomography, gold nanoparticles imaging, fast 3D acquisition, quantitative image analysis

## Abstract

Delivery of gold nanoparticles (GNPs) into live cells has high potentials, ranging from molecular-specific imaging, photodiagnostics, to photothermal therapy. However, studying the long-term dynamics of cells with GNPs using conventional fluorescence techniques suffers from phototoxicity and photobleaching. Here, we present a method for 3-D imaging of GNPs inside live cells exploiting refractive index (RI) as imaging contrast. Employing optical diffraction tomography, 3-D RI tomograms of live cells with GNPs are precisely measured for an extended period with sub-micrometer resolution. The locations and contents of GNPs in live cells are precisely addressed and quantified due to their distinctly high RI values, which was validated by confocal fluorescence imaging of fluorescent dye conjugated GNPs. In addition, we perform quantitative imaging analysis including the segmentations of GNPs in the cytosol, the volume distributions of aggregated GNPs, and the temporal evolution of GNPs contents in HeLa and 4T1 cells.

**Abbreviations:**

GNPs: gold nanoparticles
RI: refractive index
ODT: optical diffraction tomography
DMD: digital micromirror device

## 1. Introduction

Gold nanoparticles (GNPs) have been widely applied to biological cell studies in various research fields because of their distinctive properties which differentiate them from conventional biomolecules and even from other metal nanoparticles[1]. GNPs exhibit surface plasmon resonance which arises from the collective oscillation of conduction electrons by incident photons in a specific wavelength, resulting in the strong absorption of light at the specific wavelength. In addition, the size and shape of GNPs can be controlled by various synthesis techniques[1] for spectral multiplexing. Moreover, GNPs show relatively high chemical stability compared to other metal nanoparticles. With these unique properties, the current uses of GNPs can be listed from photodiagnostics to photothermal therapy[2–6].

In order to visualize GNPs inside cells and investigate the effects of GNPs, various imaging techniques have been implemented. While transmission electron microscopy (TEM) conventionally provides high spatial resolution images of GNPs, TEM requires the extremely vacuum and high-intensity electron beam, thus incompatible with live cells[7, 8]. Fluorescence imaging techniques, such as epifluorescence microscopy and confocal microscopy, have been utilized for observing the spatial distribution of GNPs inside biological cells. By tagging fluorescent dyes to the functionalized surface of GNPs, these fluorescence techniques provide high molecular specificity in live cell imaging[9, 10]. However, fluorescent probes may alter the original physiological conditions of biological cells and may also suffer from phototoxicity and photobleaching[11, 12]. In addition, three-dimensional (3-D) fluorescence imaging requires the axial scanning scheme, which suffers long acquisition time for 3-D images[13].

Holographic microscopy has been employed in imaging GNPs inside live cells, because gold has distinct refractive index (RI) values compared to cellular components, resulting in detectable optical phase shifts of the incident light. Heterodyne digital holographic microscopy was used to localize a single GNP attached to the cellular surface receptor[14]. Dark-field digital holographic microscopy was employed to track GNPs diffusing in water[15]. Wide-field interferometric phase microscopy was used to detect the photothermal phase signals from GNPs inside cells[16]. Silver nanoparticles were used to enhance molecular specificity in lensfree holography[17]. Previous techniques have measured 2-D complex optical fields, and numerical propagation of the measured optical fields employing the angular spectrum method can localize the 3-D positions of single GNPs[18]. However, these previous approaches have exhibited poor axial resolution that cannot be applied for volumetric reconstructions of complicated shaped or aggregated GNPs and cells. More importantly, 2-D complex optical fields only provide optical phase shift images, in which RI values and shapes of objects are coupled.

Recently, optical diffraction tomography (ODT) or holotomography (HT) has emerged as a label-free 3-D imaging technique, which can measure the 3-D RI distribution of biological samples in the high spatial resolution[19–21]. Since RI is an intrinsic optical property of material, ODT does not need the use of labeling agents or any additional preparations of cells. Because ODT takes into account of light diffraction inside samples and reconstructs the 3-D RI distribution of samples from measured complex optical fields, ODT can be applied for visualization of complicated structures of samples which cannot be explored by angular spectrum method[21]. Moreover, ODT provides quantitative information on biological samples from measured 3-D RI distribution, including protein concentration, cellular dry mass, and volume. Due to aforementioned advantages of ODT, it has been widely used to study various biological samples such as red blood cells[22–24], white blood cells[25, 26], neuron cells[27, 28], hepatocytes[29], bacteria[30, 31], phytoplankton[32], and human downy hair[33].

Here, we present label-free high-resolution 3-D imaging of GNPs in live cells by implementing ODT. By employing an ODT system based on a Mach-Zehnder interferometer and a digital micromirror device (DMD), the 3D RI tomograms of individual live cells with GNPs were measured. To reconstruct a 3-D RI tomogram, multiple 2-D optical field images of individual live cells were obtained with various illumination angles which were controlled by using a DMD, and then a reconstruction algorithm to inverse light scattering was applied. The measured tomograms showed that RI values of GNPs were significantly higher than other intracellular components in cells. Using this fact, GNPs were successfully segmented from cytoplasm with high spatial resolution, which was confirmed by confocal fluorescence imaging. Using the present technique, we quantitatively analyzed the volume distribution of aggregated GNPs and the time evolution of GNPs volume ratios inside two different cancer cell lines (HeLa and 4T1), which clearly provided quantitative information about the accumulation of GNPs inside cells.

## 2. Materials and Methods

### 2.1 Tomogram reconstruction

From multiple 2-D optical fields with various illumination angles, the 3-D RI distribution of a sample was reconstructed via an ODT algorithm[19, 34]. In the ODT algorithm, each 2-D Fourier spectra information corresponding to 2-D complex optical fields obtained with a specific incident angle was mapped onto a spherical surface, called Ewald sphere, in the 3-D Fourier space, according to Fourier diffraction theorem. Because an objective lens has such limited numerical aperture (NA) for collecting diffracted light from a sample, the mapped Fourier space inevitably has missing information, which is called as the missing cone. To fill the missing information, we applied Gerchberg-Papoulis algorithm with a non-negativity constraint. The details of this algorithm can be found in the previous reports[19, 33, 35]. Finally, the 3-D RI distribution of the sample was reconstructed by applying inverse 3-D Fourier transform to the mapped information in the 3-D Fourier space. More details about the principle of ODT, the ODT algorithm, and the Gerchberg-Papoulis algorithm can be found elsewhere[19, 21, 34, 36]. Reconstruction and visualization of the 3-D RI tomograms were performed using commercial software (TomoStudo, Tomocube Inc., Republic of Korea)

### 2.2 Cell culture and GNPs treatment

Human cervical cancer cells (HeLa cells) in Dulbecco's modified Eagle's medium (DMEM) and murine breast cancer cells (4T1 cells) in RPMI 1640 medium were respectively cultured at 37°C in 5% CO_2_. PEGylated and fluorescent GNPs were made from colloidal GNPs with a diameter of 20 nm. Each GNPs was added to the cell suspension at 150 μM Au ion concentration and incubated for 6 h at 37°C.

### 2.3 Fourier transform light scattering

Fourier transform light scattering (FTLS) was used to measure RI values for a GNPs solution[37, 38]. First, 2-D field images of colloidal solutions of GNPs with silica beads of known RI value (*n* = 1.4607 at *λ* = 532 nm) were taken. From the measured field image, the far-field angle-resolved light scattering spectra of silica beads were calculated by applying 2-D Fourier transformation. According to the Mie scattering theory, the angle-resolved light scattering spectra is analytically determined from the RI and size of a spherical scatterer, the RI of a medium (*n* = 1.337), and the wavelength of incident light (*λ* = 532 nm). Fitting the measured angle-resolved light scattering spectra to the Mie scattering theory provided information about the GNP solution because the other parameters were known. Calculations of the Fourier transform light scattering, and fittings to the Mie theory were performed using a custom MatLab^TM^ code.

### 2.4 Preparation of PEGylated GNPs and fluorescence-conjugated GNPs

For the preparation of PEGylated 20 nm GNPs, 20 mL of 1.0 mM HAuCl_4_ was brought to a boil in a 100 mL bottle, and then 1.6 mL of 38.8 mM tri-sodium citrate was added. The solution color changed from clear to bright red and dark red over 5 minutes. To modify the surface of GNPs with methoxy polyethylene glycol (mPEG, 5K), 25 mg of mPEG was added in bare GNPs solution and was vortexed for overnight at room temperature. After synthesis and surface modification, hydrodynamic size of the GNPs was characterized using dynamic light scattering and TEM. In order to conjugate Alexa Fluor 555 dye (Alexa Fluor 555 Succinimidyl Ester, ThermoFisher Scientific, United States) with GNPs, 20 mg of mPEG and 5 mg NH2-PEG was added to citrated GNPs and dialyzed for 2 days. 5.0 μg μL^-1^ of AF555 was added to the GNPs solution and vortexed for overnight at room temperature. Excess dyes were removed via centrifugation.

## 3. Results and Discussion

### 3.1 Principles of imaging GNPs with ODT

The schematic procedure for imaging GNPs in live cells using ODT is depicted in Fig. 1(a). Prepared PEGylated GNPs were taken up by cells before measurements (See 2.2). Live cells were placed between coverslips with a spacer and measured with an ODT system. Multiple 2-D holograms were measured with various illumination angles as shown in the subsets in Fig. 1(a). From the each measured hologram, an optical field image consisting of amplitude and phase images was retrieved by applying a field retrieval algorithm[39]. From the multiple 2-D optical field images, the 3-D RI distribution of a sample was reconstructed using the ODT algorithm based on the Fourier diffraction theorem (See 2.1)[33, 39].

**Figure 1.**
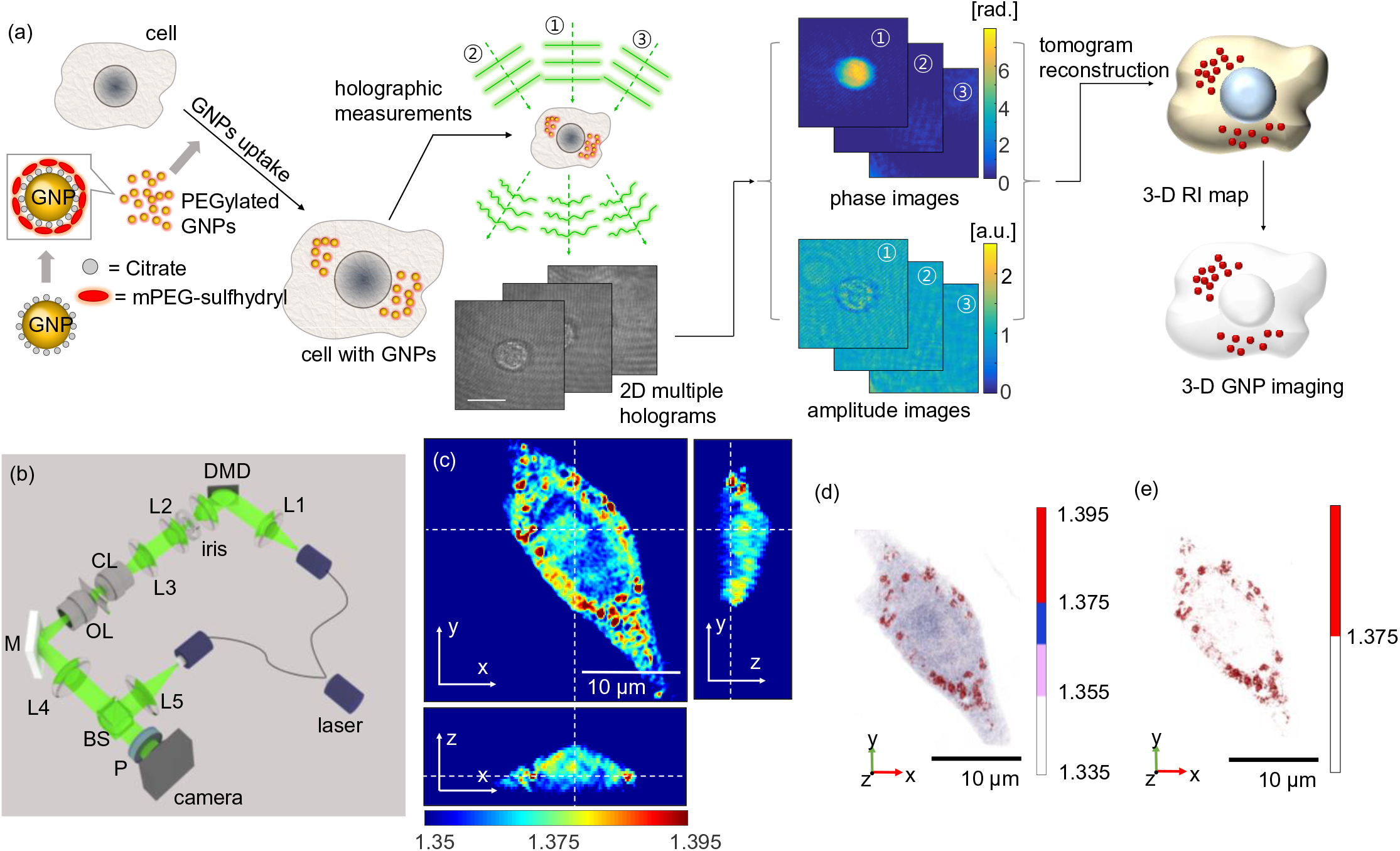
(a) Schematic procedure of imaging GNPs inside cells with ODT. **(b)** Optical setup. BS, beam splitter; DMD, digital micromirror device; OL, objective lens; CL, condenser lens; M, mirror; P, polarizer; L1-5, lenses. **(c)** Cross-sectional slices of the 3-D RI tomogram in the *x-y*, *x-z*, and *y-z* planes of a HeLa cell treated with GNPs. **(d)** The 3-D rendered image of the same cell. **(e)** The 3-D rendered image of the cell with the RI values greater than 1.375, which are considered as GNPs.

### 3.2 Experimental setup

In order to measure 3-D RI distributions of cells, we utilized commercialized ODT setups (HT-1S and HT-1H, Tomocube, Inc., South Korea). The schematic of the setup is shown in Fig. 1(b). The system was based on a Mach-Zehnder interferometric microscope equipped with a DMD. A laser beam from a diode-pumped solid-state laser (*λ* = 532 nm, 10 mW) was divided into two arms. One beam was used as a reference beam, and the other beam illuminated a sample with various incident angles ranging from −60° to 60°. In order to control the angle of incident illumination, a DMD (DLP6500FYE, Texas Instruments, United States) was implemented. By projecting a Lee hologram pattern on a DMD, the angle of the diffracted light from the DMD was precisely controlled by applying corresponding hologram patterns[40, 41]. The beam diffracted from the sample was collected by using a 60× objective lens, and then was interfered with the reference beam, generating spatially modulated holograms. The holograms of a sample were recorded with a high-speed image sensor (CMOS camera, FL3-U3-13Y3M-C, FLIR Systems, Inc., United States) with a frame rate of 300 Hz. The results in Fig. 1, Fig. 3, and Fig. 4 were obtained with an objective lens with a numerical aperture (NA) of 1.2, and the rest of results were obtained using an objective lens with an NA of 0.8. In addition, time-averaged sinusoidal pattern, a modified version of binary Lee hologram pattern, was used for the results in Fig. 3 in order to decrease the noise level[42]. The theoretical optical lateral and axial resolution of the present method were calculated as 166 nm and 1.00 μm for NA = 0.8, and 110 nm and 356 nm for NA = 1.2.

**Figure 2.**
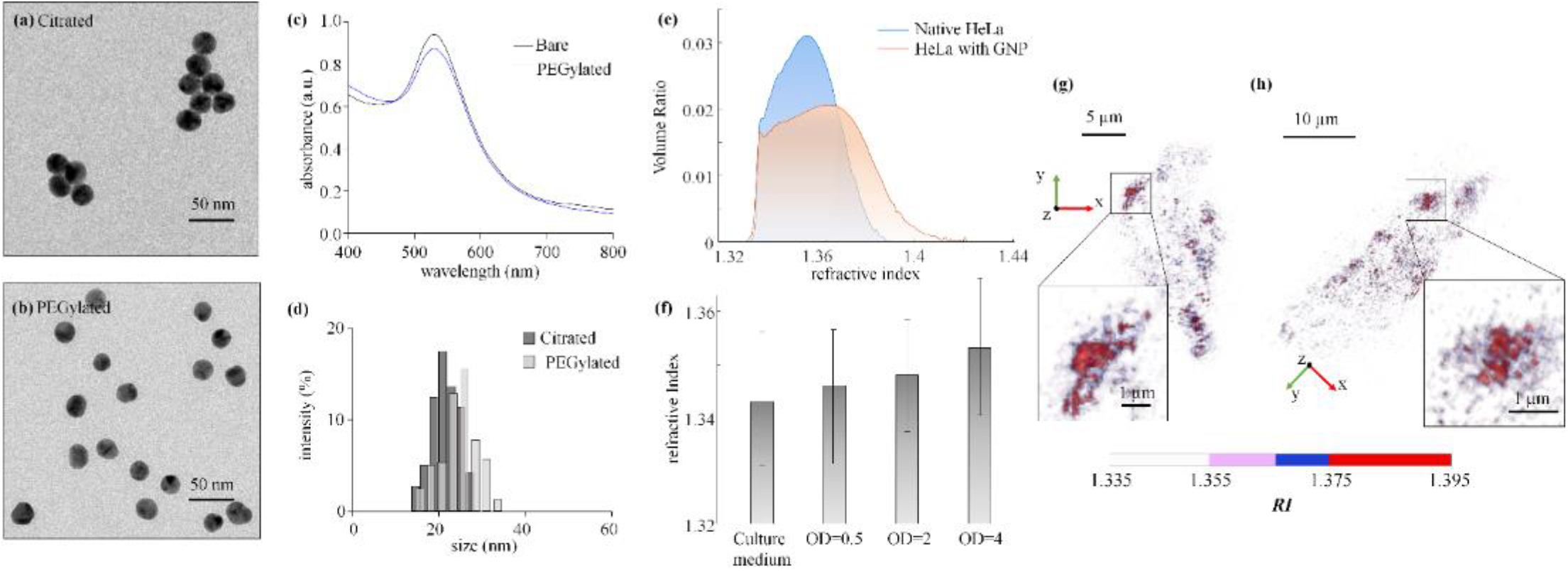
(a) Transmission electron microscopic (TEM) images of citrate and **(b)** PEGylated GNPs with average diameters of 20 nm. **(c)** Absorption spectra of citrated and PEGylated GNPs. **(d)** Hydrodynamic size distribution histogram of citrated and PEGylated GNPs. The size distribution histogram was obtained based on dynamic light scattering measurements. **(e)** Average RI histograms of GNPs treated (*n* = 15) and native cells (*n* = 15). **(f)** RI values of GNPs solutions with different concentrations of GNPs (OD values). Refractive index was measured with Fourier transform light scattering (FTLS). **(g)** and **(h)** 3-D rendered RI tomograms of aggregated GNPs.

**Figure 3.**
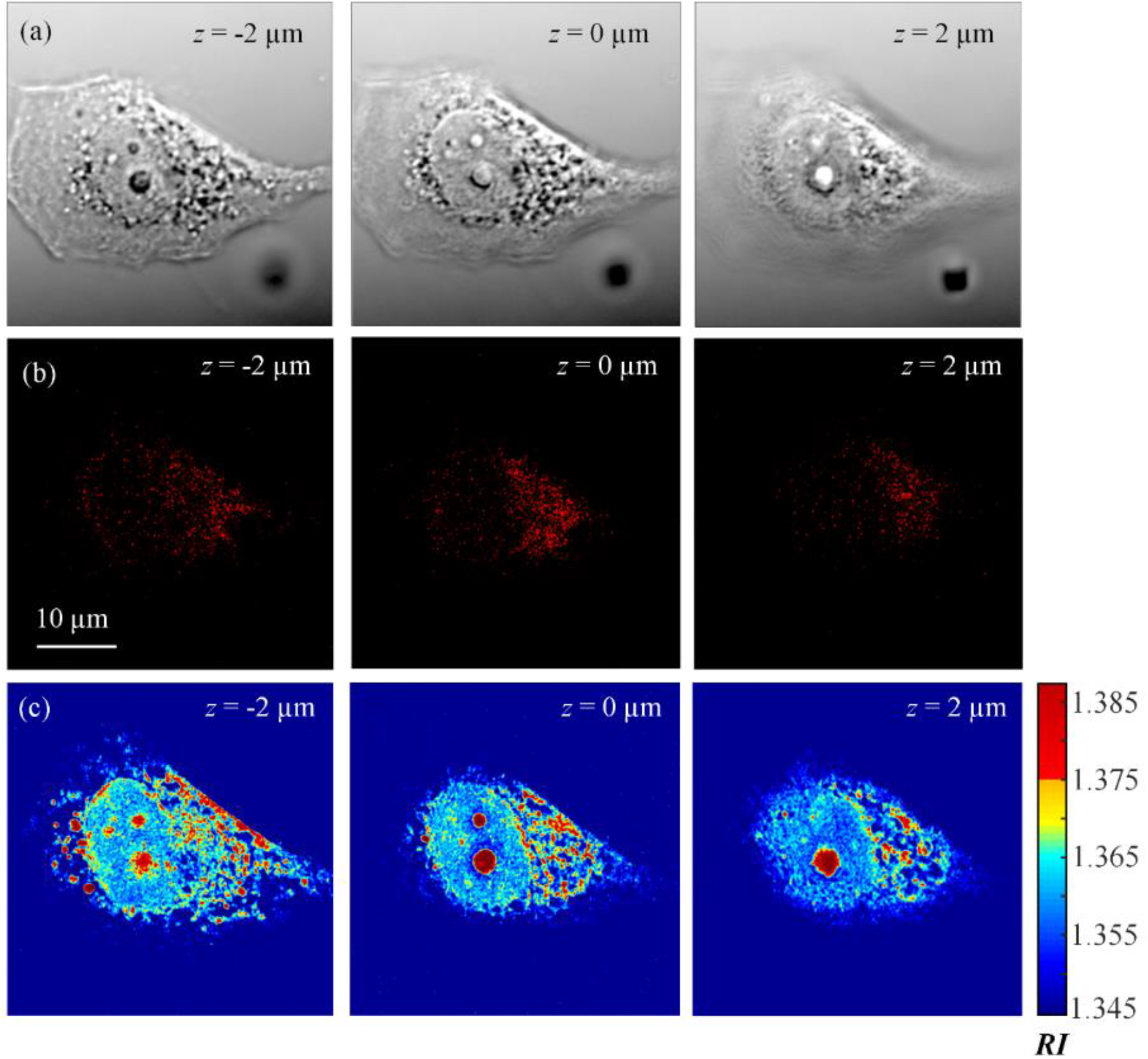
(a) Differential interference contrast (DIC) microscopic images of a HeLa cell treated with GNPs on different focal planes. **(b)** Confocal microscopic images of the same cell on same focal planes. **(c)** *x-y* cross-sections of the RI tomogram of the same cell on the same planes.

### 3.3 Synthesis and characterization of GNPs

GNPs (average diameter = 20 nm) were synthesized and further coated with methoxy-polyethyleneglycol(mPEG)-sulfhydryl (5k) molecules following the standard protocol[43]. PEGylated GNPs were stable under the experimental condition, and HeLa cells could take up individual nanoparticles. HeLa cells were treated with PEGylated GNPs solution in a cell culture medium for 6 h prior to the measurements (See 2.2).

Representative 3-D RI tomograms of GNPs inside a HeLa cell are shown in Figs. 1(c)-(e). Figure 1(c) shows cross-sectional slices of the 3-D RI tomogram in the *x-y*, *x-z*, and *y-z* planes of a HeLa cell treated with PEGylated GNPs, and 3-D rendered image of the same cell is shown in Fig. 1(d). The reconstructed 3-D RI distribution of endocytosed GNPs in the HeLa cell had regions with RI values, which were distinctively higher than those of cytoplasm. We consider these regions as the complex or aggregated GNPs in endosomes, which are highlighted in Fig. 1(e).

Synthesized GNPs were characterized by utilizing various methods [Figs. 2(a)-(d)]. First, transmission electron micrographs of citrated and PEGylated GNPs are presented in Figs. 2(a)−(b), which show their spherical shapes with uniform diameters of about 20 nm. The absorption spectra of citrated and PEGylated GNPs did not change significantly [Fig. 2(c)], indicating that no aggregation occurred during surface modification in aqueous conditions. The hydrodynamic diameters of PEGylated GNPs, which were measured using a dynamic light scattering instrument (Zetasizer Nano ZS90, Malvern Instruments Ltd., United Kingdom), were slightly larger than that of citrated GNPs because of organic molecules coated on the surface [Figure 2(d)]. The mean zeta potential of PEGylated GNPs was measured as −13 ± 1.43.

### 3.4 Comparisons between native and GNPs treated cells

In order to determine the RI distributions of GNPs inside biological cells, we measured 3-D RI tomograms of GNPs treated (*n* = 15) and native cells (*n* = 15). From the measured RI tomograms of cells, average RI histograms were calculated [Fig. 2(e)]. In order to exclude surrounding media from the calculation, voxels with RI values higher than that of surrounding media (*n* > 1.337) were only selected from the measured RI tomograms, which corresponded to cells. As shown in Fig. 2(e), GNPs treated cells showed RI distributions in which the distributions exhibited higher RI values compared to those of native cells. The maximum RI value of native cells was about 1.385, whereas that of GNPs treated cells was about 1.42. It is noteworthy that reported RI values of cell cytoplasm were within the range of 1.37-1.39. The cytoplasm, consisting of protein solutions at physiological concentrations[20, 34, 44], does not exhibit RI values higher than 1.4, except for hemozoin crystals in malaria-infected red blood cells[23] or lipid droplets in hepatocyte[29]. Therefore, these regions with RI values higher than usual cytoplasmic RI values are assumed to correspond to GNPs.

### 3.5 RI values of GNPs in colloidal and aggregated forms

To understand the high RI values measured in cells with GNPs, we measured RI values of GNPs. Due to the limited spatial resolving power of ODT, reconstructed RI tomograms were sampled with a finite voxel size of 110 × 110 × 356 nm and 166 × 166 × 1,000 nm for the high (1.2) and low (0.8) NA objective lenses, respectively. Furthermore, because of their spherical shapes, GNPs does not fill 100% of a specific voxel even for the case of externally highest concentration of colloidal or aggregations. Thus, the measured RI values should reflect the volume concentration of GNPs.

First, we measured RI values of GNPs in colloidal forms in transparent media. Various concentrations of GNPs, expressed as optical density (OD) values, were dispersed in phosphate buffered saline solution (PBS solution, pH 7.4, 50 mM, Welgene, South Korea), and the RI values of solutions were measured with Fourier transform light scattering technique[37, 38] (See 2.3). The measured RI values as a function of OD are shown in Fig. 2(f). The measured RI of the PBS solution without GNPs were 1.343 ± 0.012, which were in good agreement with the manufacturer's specification. The measured RI values for the GNP solutions increased with the concentrations of GNPs; the RI values for the GNP solutions with ODs of 0.5, 2, and 4 were 1.346 ± 0.012, 1.348 ± 0.010, and 1.353 ± 0.012, respectively. Although the measured RI values increased as the concentrations of GNPs, even the average RI values of the densest solution (OD = 4) was 1.353. However, the 3-D RI tomograms show that the presence of GNPs inside cells causes the regions with RIs higher than 1.375 [Fig. 2(e)]. Furthermore, it is physiologically unlikely that GNPs in live cells would have the concentration of higher than OD of 4.

Because the colloidal solutions of GNPs did not exhibit high RI values which we have been observed in GNPs treated cells, we hypothesized that the GNPs in cells would be in the aggregated form. To test this hypothesis, we added NaCl to colloidal GNPs solution in order to trigger the aggregation of GNPs, and measured 3-D RI tomograms of the GNP solutions with NaCl. As shown in Figs. 2(g)−(h), we observed highly aggregated parts which showed high RI values (*n* > 1.375). These high values of RI correspond to those appeared in the GNPs treated cells. These results suggest that portions of GNPs exist in a highly aggregated form, which can be imaged and identified by measuring 3-D RI tomograms of the GNP-treated cells.

Although the refractive index of gold for 532 nm is about 0.4, RI values of aggregated GNPs (*n* > 1.375) can be explained when we treat them as a mixture of gold and water. According to the Landauer-Bruggeman effective medium approximation[45], the effective RI of a mixture of gold and water exceeds 1.375 when the volumetric fraction of gold is between 0.010 and 0.190 (see Supplementary Information).

### 3.6 Correlation with confocal fluorescent images

To confirm that the regions having higher RI values inside cells represent GNPs, we compared RI tomograms of HeLa cells having PEGylated GNPs with confocal fluorescent images of the same cells. For the confocal imaging, PEGylated GNPs were further conjugated with Alexa Fluor 555 dyes. After the fluorescent GNPs treatment, cells were fixed by using 4% paraformaldehyde to prevent further movements of GNPs inside cells. Then, intracellular distributions of GNPs were visualized using a laser scanning confocal microscope (LSCM, C2 +. Nikon Instruments Inc., Japan). In order to acquire 3-D fluorescent images, samples were scanned along the axial direction at 30 different focal planes with the axial step of 0.70 μm. Total acquisition time for scanning all 30 planes was 30 min. Figures 3(a)-(c) show the representative images of a HeLa cell having PEGylated fluorescent GNPs measured by a differential interference contrast (DIC) microscope, LSCM, and ODT, respectively. Voxels with RI values higher than 1.375 show strong spatial correlation with fluorescent signals of confocal images, notwithstanding some nonmatching regions. Nevertheless, such nonmatching regions do not possess significant portions and voxels with RI values higher than 1.375 take 0.87% of total volume of native cells as calculated from RI histograms, supporting that the standard of *n* = 1.375 can be used in the imaging of GNPs with ODT.

### 3.7 Quantitative imaging analysis

Quantitative analysis for the size of aggregated GNPs inside cells has been carried out. First, the total number of aggregated GNPs was counted, and their volume ratio to total cellular volume was calculated for the sample represented in Figs. 4(a)-(c). Among the voxels with RI values higher than *n* > 1.375, a group of connected voxels was considered as one aggregated GNPs, and the volume of each group was calculated. In order to minimize the effect of noises, we excluded groups with size less than three voxels from the calculation. The total number of aggregated GNPs was 150, and the volume fraction of those GNPs to the cellular volume (1.57×10^3^ μm^3^) was 18.31%. Furthermore, the volume distribution of aggregated GNPs was organized and compared with the results for 10 different samples as shown in Fig. 4(d). Groups of aggregated GNPs with a volume smaller than 0.1 μm^3^ were considered as the ‘weakly aggregated GNPs,’ and those with bigger volume were considered as the ‘highly aggregated GNPs.’ The distribution of weakly aggregated GNPs showed a non-Gaussian distribution that the number of groups continuously decreased as the volume increased. The average number of highly aggregated GNPs for 10 cells was 16.4 ± 15.6.

**Figure 4.**
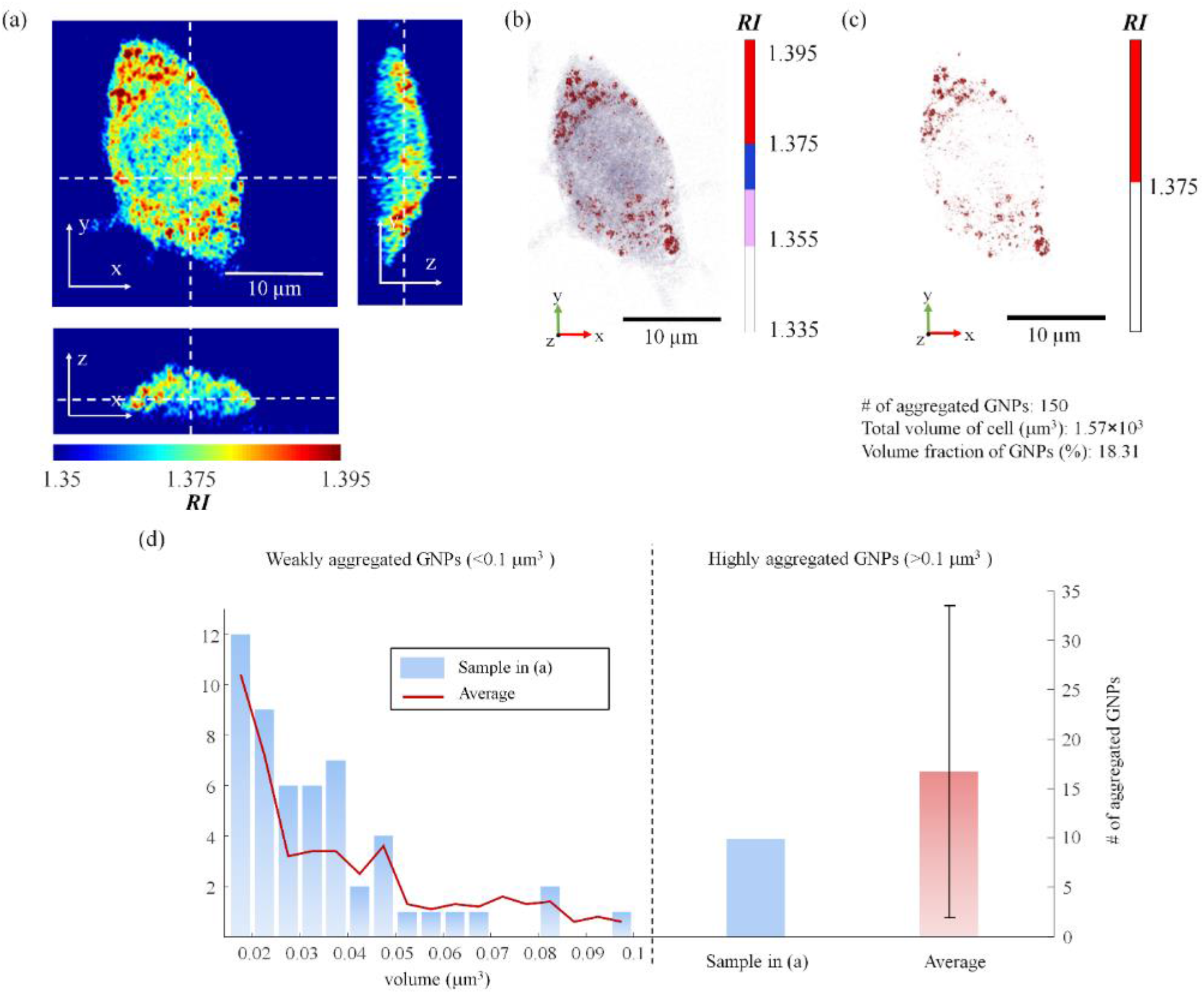
(a) Cross-sectional slices of the 3-D RI tomogram in the *x-y*, *x-z*, and *y-z* planes of a HeLa cell treated with GNPs. **(b)** The 3-D rendered image of the same cell. **(c)** The 3-D rendered image of the cell with the RI values greater than 1.375, which are considered as GNPs. **(d)** Volume distribution of aggregated GNPs inside HeLa cells.

Then, we quantitatively measured the temporal evolution of volumetric ratios of GNPs inside HeLa cells during the GNPs treatment times (0 min, 5 min, 1 h, 3 h, and 6 h). RI tomograms of 10 cells for each treatment time were acquired as shown in Fig. 5(a), from which cellular volume and the volume of GNPs were calculated by integrating voxels with RI values higher than *n* > 1.337 and *n* > 1.375, respectively. The volume proportion of GNPs inside HeLa cells increased from 2.45% at 0 min to 31.01% after 6 h of GNPs treatment [Fig. 5(b)]. The results imply that GNPs are continuously transported into HeLa cells, which is in good agreement with previous reports[46–48]. This result was also confirmed by another mammalian cell line (4T1 cell, murine breast cancer cell) as the volume proportion of GNPs inside 4T1 cells (*n* = 10) increased from 2.15% at 0 min to 46.0% after 6 h of GNPs treatment [Figs. 5(c)-(d)]. These results demonstrate the ability of ODT as a tool for the quantitative 3-D analysis of nanoparticles because the axial scanned 2-D image set of confocal microscopy does not provide the information for axial connectivity between particles as well as the size of resulted point spread function can be varied along the gain and laser intensity.

**Figure 5.**
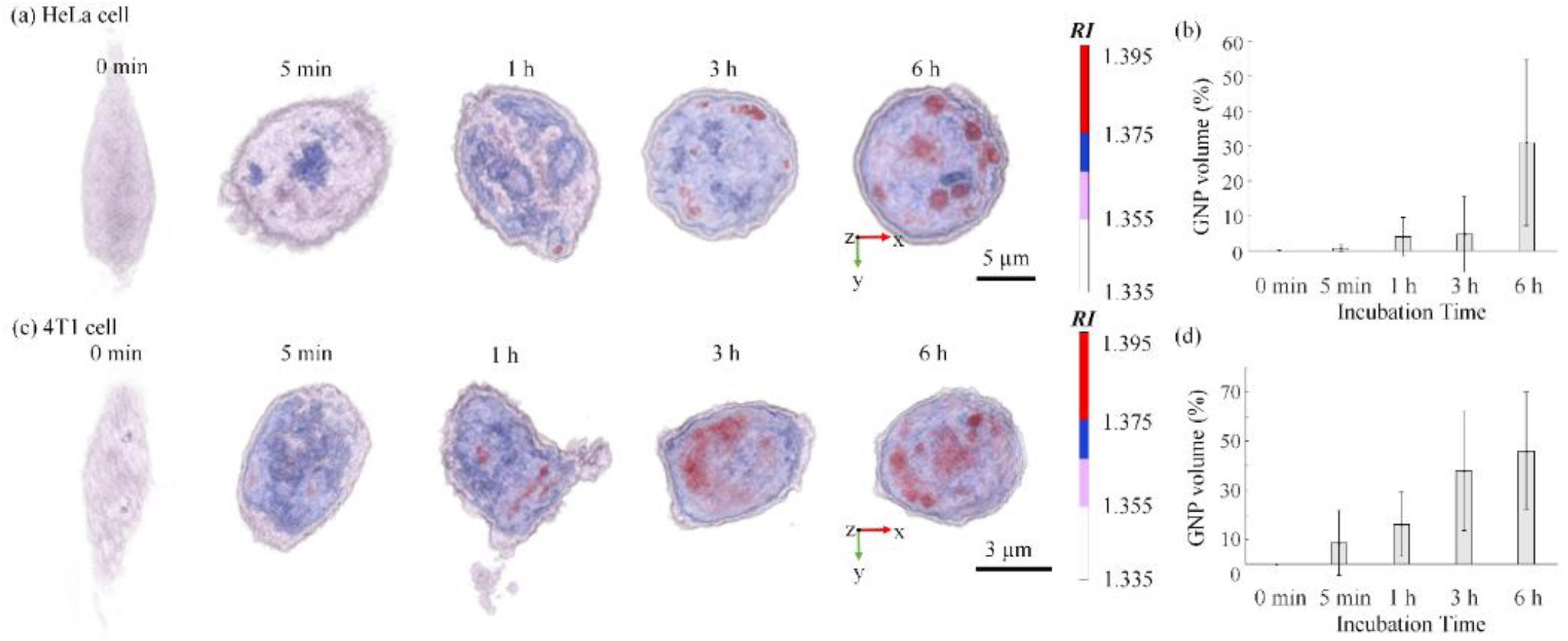
(a) 3-D rendered RI tomograms of representative HeLa cells with different GNPs treatment time. **(b)** The average volume ratio of GNPs in HeLa cells along the treatment time. **(c)** 3-D rendered RI tomograms of representative 4T1 cells with different GNPs treatment times. **(d)** The average volume ratio of GNPs in 4T1 cells along the treatment time.

## 4 Concluding remarks

We present a label-free 3-D imaging method for measuring the 3-D spatial distribution of GNPs inside live cells by employing ODT. The 3-D RI distribution of GNPs inside live cells reconstructed via ODT shows significantly high RI values (*n* > 1.375) compares to surrounding cytoplasm, which was also confirmed by fluorescence images of the same GNPs. This fact was applied for the segmentation of the spatial distribution of GNPs from measured RI tomograms of cells. In addition, the present method provides quantitative analysis of the volume distribution of aggregated GNPs and time evolution of the transport of GNPs inside various types of mammalian cell lines.

Possible errors can come out from the use of specific cells containing cellular materials with high RI values comparable to GNPs. For example, lipid droplets and carbohydrates inside cells have RI values overlapped with those of GNPs, which means present method cannot resolve them from GNPs inside cells. In addition, the sensitivity of present method should be increased in order to visualize GNPs in colloidal and weakly aggregated forms. It is expected that the employment of hyperspectral ODT[49, 50] or external photothermal excitation of GNPs[51] can resolve this problem by obtaining the dispersion of GNPs for a different wavelength or by detecting photothermal signals from GNPs which are expressed as the changes in phase delay maps.

We expect that the present technique can be utilized for cancer cell imaging by the conjugation of GNPs with tumour binding antibodies[52]. Moreover, the image stitching method can expand the field of view making tissue imaging suitable [53], as well as this method, can be employed in the long-term tracking of GNPs in tissues by using tissue phase imaging techniques and compact live cell incubators. Furthermore, recent developments of beam scanning devices and detectors in ODT enable the fast acquisition of the 3-D RI distribution of samples with the tomogram acquisition rate of 100 Hz[40, 54, 55]. Recently, several commercial ODT instruments were released, and thus, lowering barriers to entry for the active use of ODT in research areas such as biomedical imaging and disease diagnostics. In a sense, this work shows not only the new GNPs imaging method but also one of the possible biomedical applications of ODT.

## 5 Funding

This work was supported by KAIST, Tomocube, and the National Research Foundation of Korea (2015R1A3A2066550, 2014M3C1A3052567, 2014K1A3A1A09063027 and 2015R1A1A1A05001420), and the National Research Council of Science & Technology (NST) grant (CAP-14-03-KRISS) funded by the Ministry of Science, ICT & Future Planning, Republic of Korea.

## Conflicts of Interest

Prof. Y.K.P and Dr. K.H. have financial interests in Tomocube Inc., a company that commercializes optical diffraction tomography and quantitative phase imaging instruments and is one of the sponsors of the work.

## References

[1] S.K. Ghosh, T. Pal, Interparticle coupling effect on the surface plasmon resonance of gold nanoparticles: from theory to applications, Chemical reviews 107(11) (2007) 4797–4862.

[2] S.K. Balasubramanian, L. Yang, L.-Y.L. Yung, C.-N. Ong, W.-Y. Ong, E.Y. Liya, Characterization, purification, and stability of gold nanoparticles, Biomaterials 31(34) (2010) 9023–9030.

[3] M. Grzelczak, J. Pérez-Juste, P. Mulvaney, L.M. Liz-Marzán, Shape control in gold nanoparticle synthesis, Chemical Society Reviews 37(9) (2008) 1783–1791.

[4] N.J. Durr, T. Larson, D.K. Smith, B.A. Korgel, K. Sokolov, A. Ben-Yakar, Two-photon luminescence imaging of cancer cells using molecularly targeted gold nanorods, Nano Lett 7(4) (2007) 941–945.

[5] I.H. El-Sayed, X. Huang, M.A. El-Sayed, Selective laser photo-thermal therapy of epithelial carcinoma using anti-EGFR antibody conjugated gold nanoparticles, Cancer letters 239(1) (2006) 129–135.

[6] X. Huang, I.H. El-Sayed, W. Qian, M.A. El-Sayed, Cancer cell imaging and photothermal therapy in the near-infrared region by using gold nanorods, Journal of the American Chemical Society 128(6) (2006) 2115–2120.

[7] X. Huang, P.K. Jain, I.H. El-Sayed, M.A. El-Sayed, Gold nanoparticles: interesting optical properties and recent applications in cancer diagnostics and therapy, (2007).

[8] C.M. Pitsillides, E.K. Joe, X. Wei, R.R. Anderson, C.P. Lin, Selective cell targeting with light-absorbing microparticles and nanoparticles, Biophysical journal 84(6) (2003) 4023–4032.

[9] B.D. Chithrani, A.A. Ghazani, W.C. Chan, Determining the size and shape dependence of gold nanoparticle uptake into mammalian cells, Nano Lett 6(4) (2006) 662–668.

[10] L.F. Kourkoutis, J.M. Plitzko, W. Baumeister, Electron microscopy of biological materials at the nanometer scale, Annual review of materials research 42 (2012) 33–58.

[11] F. Sousa, S. Mandal, C. Garrovo, A. Astolfo, A. Bonifacio, D. Latawiec, R.H. Menk, F. Arfelli, S. Huewel, G. Legname, Functionalized gold nanoparticles: a detailed in vivo multimodal microscopic brain distribution study, Nanoscale 2(12) (2010) 2826–2834.

[12] Y. Zhang, M. Hensel, Evaluation of nanoparticles as endocytic tracers in cellular microbiology, Nanoscale 5(19) (2013) 9296–9309.

[13] Y. Fei, Y.-S. Sun, Y. Li, K. Lau, H. Yu, H.A. Chokhawala, S. Huang, J.P. Landry, X. Chen, X. Zhu, Fluorescent labeling agents change binding profiles of glycan-binding proteins, Molecular BioSystems 7(12) (2011) 3343–3352.

[14] R. Hoebe, C. Van Oven, T.J. Gadella, P. Dhonukshe, C. Van Noorden, E. Manders, Controlled light-exposure microscopy reduces photobleaching and phototoxicity in fluorescence live-cell imaging, Nature biotechnology 25(2) (2007) 249–253.

[15] F. Verpillat, F. Joud, P. Desbiolles, M. Gross, Dark-field digital holographic microscopy for 3D-tracking of gold nanoparticles, Optics express 19(27) (2011) 26044–26055.

[16] L. Shang, R.M. Dörlich, S. Brandholt, R. Schneider, V. Trouillet, M. Bruns, D. Gerthsen, G.U. Nienhaus, Facile preparation of water-soluble fluorescent gold nanoclusters for cellular imaging applications, Nanoscale 3(5) (2011) 2009–2014.

[17] Q. Wei, E. McLeod, H. Qi, Z. Wan, R. Sun, A. Ozcan, On-chip cytometry using plasmonic nanoparticle enhanced lensfree holography, Scientific reports 3 (2013) 1699.

[18] N. Warnasooriya, F. Joud, P. Bun, G. Tessier, M. Coppey-Moisan, P. Desbiolles, M. Atlan, M. Abboud, M. Gross, Imaging gold nanoparticles in living cell environments using heterodyne digital holographic microscopy, Optics Express 18(4) (2010) 3264–3273.

[19] E. Wolf, Three-dimensional structure determination of semi-transparent objects from holographic data, Optics Communications 1(4) (1969) 153–156.

[20] K. Lee, K. Kim, J. Jung, J.H. Heo, S. Cho, S. Lee, G. Chang, Y.J. Jo, H. Park, Y.K. Park, Quantitative phase imaging techniques for the study of cell pathophysiology: from principles to applications, Sensors 13(4) (2013) 4170–4191.

[21] K. Kim, H.-O. Yoon, M. Diez-Silva, M. Dao, R. Dasari, Y.-K. Park, High-resolution three-dimensional imaging of red blood cells parasitized by *Plasmodium falciparum* and *in situ* hemozoin crystals using optical diffraction tomography, J. Biomed. Opt. 19(1) (2014) 011005–12.

[22] H. Park, S.-H. Hong, K. Kim, S.-H. Cho, W.-J. Lee, Y. Kim, S.-E. Lee, Y. Park, Characterizations of individual mouse red blood cells parasitized by Babesia microti using 3-D holographic microscopy, Scientific Reports 5 (2015).

[23] K. Kim, K.S. Kim, H. Park, J.C. Ye, Y. Park, Real-time visualization of 3-D dynamic microscopic objects using optical diffraction tomography, Optics Express 21(26) (2013) 32269–32278.

[24] H. Park, T. Ahn, K. Kim, S. Lee, S.-y. Kook, D. Lee, I.B. Suh, S. Na, Y. Park, Three-dimensional refractive index tomograms and deformability of individual human red blood cells from cord blood of newborn infants and maternal blood, Journal of biomedical optics 20(11) (2015) 111208–111208.

[25] J. Yoon, K. Kim, H. Park, C. Choi, S. Jang, Y. Park, Label-free characterization of white blood cells by measuring 3D refractive index maps, Biomed Opt Express 6(10) (2015) 3865–3875.

[26] J. Yoon, Y. Jo, M.-h. Kim, K. Kim, S. Lee, S.-J. Kang, Y. Park, Label-free identification of non-activated lymphocytes using three-dimensional refractive index tomography and machine learning, bioRxiv (2017) 107805.

[27] S.-A. Yang, J. Yoon, K. Kim, Y. Park, Measurements of morphological and biochemical alterations in individual neuron cells associated with early neurotoxic effects in Parkinson's disease using optical diffraction tomography, Cytometry part A (2017).

[28] Y. Cotte, F. Toy, P. Jourdain, N. Pavillon, D. Boss, P. Magistretti, P. Marquet, C. Depeursinge, Marker-free phase nanoscopy, Nature Photonics 7(2) (2013) 113–117.

[29] K. Kim, S. Lee, J. Yoon, J. Heo, C. Choi, Y. Park, Three-dimensional label-free imaging and quantification of lipid droplets in live hepatocytes, Scientific reports 6 (2016) 36815.

[30] T.I. Kim, B. Kwon, J. Yoon, I.-J. Park, G.S. Bang, Y. Park, Y.-S. Seo, S.-Y. Choi, Antibacterial Activities of Graphene Oxide-Molybdenum Disulfide Nanocomposite Films, ACS Applied Materials & Interfaces (2017).

[31] M. Bennet, D. Gur, J. Yoon, Y. Park, D. Faivre, A Bacteria-Based Remotely Tunable Photonic Device, Advanced Optical Materials (2016).

[32] S. Lee, K. Kim, A. Mubarok, A. Panduwirawan, K. Lee, S. Lee, H. Park, Y. Park, High-Resolution 3-D Refractive Index Tomography and 2-D Synthetic Aperture Imaging of Live Phytoplankton, Journal of the Optical Society of Korea 18(6) (2014) 691–697.

[33] K. Kim, H. Yoon, M. Diez-Silva, M. Dao, R.R. Dasari, Y. Park, High-resolution three-dimensional imaging of red blood cells parasitized by Plasmodium falciparum and in situ hemozoin crystals using optical diffraction tomography, Journal of biomedical optics 19(1) (2014) 011005–011005.

[34] K. Kim, J. Yoon, S. Shin, S. Lee, S.-A. Yang, Y. Park, Optical diffraction tomography techniques for the study of cell pathophysiology, Journal of Biomedical Photonics & Engineering 2(2) (2016) 020201.

[35] C.R. Patra, R. Bhattacharya, D. Mukhopadhyay, P. Mukherjee, Fabrication of gold nanoparticles for targeted therapy in pancreatic cancer, Advanced drug delivery reviews 62(3) (2010) 346–361.

[36] J. Lim, K. Lee, K.H. Jin, S. Shin, S. Lee, Y. Park, J.C. Ye, Comparative study of iterative reconstruction algorithms for missing cone problems in optical diffraction tomography, Optics Express 23(13) (2015) 16933–16948.

[37] H. Ding, Z. Wang, F. Nguyen, S.A. Boppart, G. Popescu, Fourier transform light scattering of inhomogeneous and dynamic structures, Physical review letters 101(23) (2008) 238102.

[38] J.H. Jung, J. Jang, Y. Park, Spectro-refractometry of individual microscopic objects using swept-source quantitative phase imaging, Analytical chemistry 85(21) (2013) 10519–25.

[39] S.K. Debnath, Y. Park, Real-time quantitative phase imaging with a spatial phase-shifting algorithm, Optics Letters 36(23) (2011) 4677–4679.

[40] S. Shin, K. Kim, J. Yoon, Y. Park, Active illumination using a digital micromirror device for quantitative phase imaging, Optics Letters 40(22) (2015) 5407–5410.

[41] S. Shin, K. Kim, T. Kim, J. Yoon, K. Hong, J. Park, Y. Park, Optical diffraction tomography using a digital micromirror device for stable measurements of 4-D refractive index tomography of cells, Proc. of SPIE 9718 (2016) 971814.

[42] K. Lee, K. Kim, G. Kim, S. Shin, Y. Park, Time-multiplexed structured illumination using a DMD for optical diffraction tomography, arXiv preprint arXiv:1612.00044 (2016).

[43] K. Lee, K. Kim, J. Jung, J. Heo, S. Cho, S. Lee, G. Chang, Y. Jo, H. Park, Y. Park, Quantitative phase imaging techniques for the study of cell pathophysiology: from principles to applications, Sensors 13(4) (2013) 4170–4191.

[44] G. Popescu, Quantitative Phase Imaging of Cells and Tissues, McGraw-Hill Professional 2011.

[45] D. Stroud, The effective medium approximations: Some recent developments, Superlattices and microstructures 23(3) (1998) 567–573.

[46] K. Kim, Y. Park, Fourier transform light scattering angular spectroscopy using digital inline holography, Optics letters 37(19) (2012) 4161–4163.

[47] H. Yu, H. Park, Y. Kim, M.W. Kim, Y. Park, Fourier-transform light scattering of individual colloidal clusters, Optics Letters 37(13) (2012) 2577–2579.

[48] K. Song, P. Xu, Y. Meng, F. Geng, J. Li, Z. Li, J. Xing, J. Chen, B. Kong, Smart gold nanoparticles enhance killing effect on cancer cells, International journal of oncology 42(2) (2013) 597–608.

[49] H.C. Lin, H.H. Lin, C.Y. Kao, A.L. Yu, W.P. Peng, C.H. Chen, Quantitative Measurement of Nano-/Microparticle Endocytosis by Cell Mass Spectrometry, Angewandte Chemie International Edition 49(20) (2010) 3460–3464.

[50] O. Betzer, R. Meir, T. Dreifuss, K. Shamalov, M. Motiei, A. Shwartz, K. Baranes, C.J. Cohen, N. Shraga-Heled, R. Ofir, In-vitro Optimization of Nanoparticle-Cell Labeling Protocols for In-vivo Cell Tracking Applications, Scientific reports 5 (2015).

[51] J. Jung, K. Kim, H. Yu, K. Lee, S. Lee, S. Nahm, H. Park, Y. Park, Biomedical applications of holographic microspectroscopy [Invited], Applied Optics 53(27) (2014) G111–G122.

[52] J.-H. Jung, J. Jang, Y. Park, Spectro-refractometry of individual microscopic objects using swept-source quantitative phase imaging, Analytical chemistry 85(21) (2013) 10519–10525.

[53] N.A. Turko, A. Peled, N.T. Shaked, Wide-field interferometric phase microscopy with molecular specificity using plasmonic nanoparticles, Journal of biomedical optics 18(11) (2013) 111414–111414.

[54] Y. Sung, W. Choi, C. Fang-Yen, K. Badizadegan, R.R. Dasari, M.S. Feld, Optical diffraction tomography for high resolution live cell imaging, Optics express 17(1) (2009) 266–277.

[55] K. Kim, J. Yoon, Y. Park, Simultaneous 3D visualization and position tracking of optically trapped particles using optical diffraction tomography, Optica 2(4) (2015) 343–346.

